# “Dark taxonomy”: a new protocol for overcoming the taxonomic impediments for dark taxa and broadening the taxon base for biodiversity assessment

**DOI:** 10.1101/2023.08.31.555664

**Authors:** Rudolf Meier, Amrita Srivathsan, Sarah Siqueira Oliveira, Maria Isabel P.A. Balbi, Yuchen Ang, Darren Yeo, Jostein Kjærandsen, Dalton de Souza Amorim

## Abstract

We are entering the 6^th^ mass extinction with little data for “dark taxa” although they comprise most species. Much of the neglect is due to the fact that conventional taxonomic methods struggle with handling thousands of specimens belonging to hundreds of species. We thus here propose a new strategy that we call “dark taxonomy.” It addresses (1) taxonomic impediments, (2) lack of biodiversity baselines, (3) and low impact of revisionary research. Taxonomic impediments are reduced by carrying out revisions at small geographic scales to keep the number of specimens low. The risk of taxonomic error is kept low by delimiting species based on two types of data. We then show that dark taxonomy can yield important biodiversity baseline data because it uses samples obtained with biomonitoring traps. Lastly, we argue that the impact of research can be improved by publishing two manuscripts addressing different readerships. The principles of dark taxonomy are illustrated by our taxonomic treatment of Singapore’s fungus gnats (Mycetophilidae) based only on Malaise trap samples. We show that a first batch of specimens (N=1,454) contains 120 species, of which 115 are new to science thus reducing taxonomic impediments by increasing the number of described Oriental species by 25%. Species delimitation started with using DNA barcodes to estimate the number of MOTUs before “LIT” (Large-scale Integrative Taxonomy) was used to obtain the species boundaries for the 120 species by integrating morphological and molecular data. To test the taxonomic completeness of the revision, we then analyzed a second batch of 1,493 specimens and found that >97% belonged to the 120 species described based on the first batch. Indeed, the second batch only contained 18 new and rare MOTUs; i.e. our study suggests that a single revision can simultaneously yield the names for all important species and relevant biodiversity baseline data. Overall, we believe that “dark taxonomy” can quickly ready a large unknown taxon for biomonitoring.

## Introduction

Biodiversity assessment and monitoring are among the biggest and most urgent challenges of modern biology, given that many natural environments are disappearing fast and biodiversity loss is destabilizing whole ecosystems. However, due to taxonomic impediments, biodiversity assessment is difficult for many arthropod clades (e.g., Wheeler et al. 2004; Evenhuis 2007; de Carvalho et al. 2007). The problem is particularly severe in the tropics and for taxa characterized by high species diversity and abundance. For such taxa, conventional approaches to taxonomy involving morphospecies sorting tend to be ineffective, because they were not designed with hyperabundant and hyperdiverse taxa in mind (Hartop et al. 2022). These taxa are nowadays called “dark taxa,” a term that Hartop et al. (2022) suggested should be restricted to clades where the undescribed fauna is estimated to exceed the described fauna by at least one order of magnitude and the total diversity exceeds 1,000 species worldwide. Taxonomists have long learned to avoid working on such dark taxa and instead tend to work on clades with moderate numbers of specimens and species ideally in taxa that are overall already reasonably well known (i.e., with prior revisions, identification keys, illustrations etc.). Projects that satisfied these criteria have been readily available and selecting them have been good career advice for young taxonomists. However, this also meant that many truly species-rich and abundant taxa remained neglected (Hartop et al. 2022; Srivathsan et al. 2023). Arguably, this has created a special kind of taxonomic impediment (“dark taxon impediment”: Meier et al. 2022). Addressing this impediment has become high priority because taxonomic chauvinism (Bonnet et al. 2002; Troudet et al. 2017) needs to be overcome in an era where we need quantitative data on all of biodiversity.

What is needed are new taxonomic techniques that are scalable and thus suitable for dark taxa. Some examples illustrate how scalable they have to be: a recent study barcoded 7,059 specimens of scuttle flies (Diptera: Phoridae) collected by one Malaise trap over only eight weeks in a Ugandan National Park and found evidence for >650 species (Srivathsan et al. 2019). Yet only 462 scuttle fly species have been described for the entire Afrotropical region (Phorid Catalog, 2023: accessed 13 August 2023). Similarly, the diversity of the gall midges (Diptera: Cecidomyiidae) has been estimated to be 800+ species at a single site of Costa Rica (Brown et al. 2018, Borkent et al. 2018), while Hebert et al. (2016) estimated the gall midge diversity in Canada to be close to 8,500 species based on barcodes , despite fewer than 250 Nearctic species having been described for this family (Savage et al. 2019). These cases illustrate why truly hyperdiverse taxa are sometimes called “open-ended” although the term dark taxon (Hausmann et al. 2020, Hartop et al. 2022; Page 2011, 2016) is more commonly used these days. These taxa will frustrate attempts at carrying out traditional comprehensive taxonomic revisions at reasonable geographic scales (Bickel 2009) and arguably need a different set of taxonomic protocols, which we here propose to call “dark taxonomy” to highlight that these techniques are fundamentally different from what is practiced for less diverse/abundant taxa.

The first key principle of “dark taxonomy” is embracing “faunistics”, i.e., determining the geographic scope of the taxonomic treatment by restricting it to the number of specimens and species that can be reasonably handled. This is different from the traditional approach revising at a geographic scope that matches the putative range of species. Implicitly, almost all “turbo-taxonomy” studies have embraced such faunistic sampling without much discussion (e.g., Fernandez-Triana et al. 2014, 2022; Marsh et al., 2013; Riedel et al., 2013a,b; Riedel and Narakusumo 2019, but see Dijkstra et al. 2015), presumably because it goes against the conventional recommendation that taxonomic revisions should cover all known species/specimens of a clade at a geographic scale appropriate for the species included in the clade. Such “completism” is desirable, but also contributes to taxonomic chauvinism because it cannot be applied to dark taxa. Two ways to relax the requirements is to limit taxonomic treatments to a small geographic area (“faunistics”) and/or to only work on a subset of the available specimens. The taxonomic treatment underlying this study is a particularly extreme example for relaxing both requirements. We delimit, identify, and describe the species of Mycetophilidae (Diptera; fungus gnats) found only in recently collected Malaise trap samples obtained only from on a small island in the Oriental region (Singapore, 730 km^2^).

Faunistic sampling carries obvious risks: the most important one is incorrect species delimitation because intraspecific genetic variability is poorly assessed for wide-spread species (Bergsten et al. 2012). However, we predict that this error will be manageable as long as the principles of integrative taxonomy are applied; i.e., species delimitation based on multiple data sources (Dayrat 2005). It is likely that avoiding such errors will be particularly effective if the characters used for species delimitation were generated by different evolutionary processes. With regard to morphology, taxonomists usually collect information from a broad array of structures including genitalia, legs, wings etc. Genitalia are likely to be under strong sexual selection (e.g., Eberhard 1985, Rowe and Arnqvist 2012), while many other body parts are presumably more shaped by viability selection. It thus appears probable that most speciation events will leave traces in such a diverse array of structures, regardless of whether speciation was fast or slow. Of course, there are exceptions as is well known from the literature on “cryptic species” (Bickford et al. 2007). Therefore, morphology should be paired with a type of data that evolves largely via drift and thus mostly reflects time of divergence between closely related species. This is the case for DNA barcodes, given that closely related species mostly differ with regard to synonymous sites (Kwong et al. 2012; Pentinsaari et al. 2016). Large DNA barcode distances within morphospecies can therefore flag cases where morphological differentiation is lower than expected based on time of divergence. This can then trigger more detailed work to resolve species limits for taxa that have high intra- or low interspecific variability (“reciprocal illumination”). We have repeatedly found this to be very effective for resolving taxonomic problems for a range of fly families (Ang et al. 2013, Tan et al. 2010, Rohner et al. 2014, Laamanen et al. 2003).

The second element for tackling large numbers of specimens is what we call the “reverse workflow.” Traditionally, species discovery has started with sorting specimens to species using morphology. Subsequently, the morphospecies were validated with few barcodes from individuals representing the morphospecies. This workflow is ineffective for dark taxa, where even samples collected at small geographic scales routinely contain too many specimens and species for morphospecies sorting. Imagine having to sort the aforementioned >7,000 specimens of Ugandan phorids into 650 species in this manner, or the 18,000 phorids in Hartop et al. (2022) into >350 species. As argued before, such samples should first be sorted to Molecular Operational Taxonomic Units (MOTUs) with barcodes (Wang et al. 2018; Hartop et al. 2022). This is now feasible because barcodes can be acquired semi-automatically and at low cost by staff who only specialize in molecular methods (Meier et al. 2016; Srivathsan et al. 2019, 2021). Presorting large numbers of specimens to putative species means that taxonomic experts can concentrate on revising MOTU boundaries using other sources of data (Wang et al. 2018; Hartop et al. 2022).

However, delimiting large numbers of species based on two types of data still requires explicit rules for assessing and resolving conflict between data types. We recently proposed “Large-scale Integrative Taxonomy” (LIT) for this purpose (Hartop et al. 2022). The LIT protocol addresses the “integrative taxonomy conundrum” that collecting two different types of data for all specimens will slow down taxonomy when faster speed is needed to address taxonomic impediments. LIT shows that the conundrum can be overcome by fast collection of one type of data (e.g., semiautomatic acquisition of barcodes) followed by only testing MOTU boundaries by subsampling specimens for a second type of data (Hartop et al. 2022). This is what LIT implements. In the first stage, barcodes are used to generate a set of MOTUs based on one criterion (e.g., clustering at 3% pairwise distance). In the second step they are classified as “stable” or “instable” depending on whether different algorithms and clustering thresholds change MOTU composition. In the third stage, the MOTUs are tested by studying the morphology of specimens representing the main and/or most divergent haplotypes— instable MOTUs are tested more extensively than stable ones (see Hartop et al. 2022). In the fourth stage, those MOTUs that are congruent with morphological evidence are accepted as species. The fifth stage is reserved for resolving conflict between morphological and molecular evidence. As argued in Hartop et al. (2022), such conflict is predicted to be observed for recently diverged species which are expected to be incorrectly lumped into one MOTU (also due to introgression) and for “old” species with divergent allopatric populations which will be split into multiple MOTUs. Conflict can thus usually be resolved by testing whether MOTU boundaries obtained with different algorithms or clustering parameters are congruent with morphological evidence. If so, these morphospecies can also be described because the molecular and morphological data are not genuinely in conflict. If not, the MOTUs/morphospecies are left unresolved until a third data source can be used to determine species boundaries. Hartop et al. (2022) showed that only 5-10% of all specimens had to be checked to find congruent species hypotheses for the ca. 350 MOTUs that were analyzed by studying 18,000 specimens.

Sorting specimens into putative species and revising the MOTU boundaries using LIT yields species-level units. The next step is establishing which species are already described and which need description. As illustrated in our companion monograph of mycetophilids of Singapore (Amorim et al., 2023), the task of sorting through old descriptions is manageable for dark taxa in the tropics because so few species have been described. The “superficial description impediment” of Meier et al. (2022) may thus be somewhat less onerous for the taxa and regions of the planet that are most in need for dark taxonomy. In other parts of the world, revisions are more likely to get stuck after completing LIT because too many scientific names cannot be resolved. This is frustrating, but it is important to remember that delimiting species with two types of data is already a major step in the right direction. Furthermore, museomics is evolving quickly and will help with identifying which species have already been named once reliably identified or type material are sequenced (e.g., Santos et al. 2023).

Developing high-throughput taxonomic techniques for dark taxa is important, but we will also need more taxonomists to take on dark taxa. In proposing “dark taxonomy”, we suggest two ways for increasing impact and thus the interest in the taxonomy of dark taxa. The first is that initial revisions of dark taxa should be based on specimens collected with the kind of sampling techniques that are also used for monitoring biodiversity. For many groups of insects, this would be Malaise-, pitfall-, light-, and/or flight intercept traps. Revising taxa based on such samples will simultaneously yield biodiversity baselines and species descriptions that are immediately relevant for biomonitoring. Furthermore, the studies will generate DNA barcode databases that immediately become valuable for analyzing metabarcoding and metagenomics data. Of course, another benefit is also that fresh specimens are easier to process at scale, because they are suitable for rapid barcoding and all specimens have locality information in a digital format.

The second strategy is to publish the dark taxonomy studies in pairs of companion papers. The first presents results that are of interest to a broader readership such as a new approach to high throughput taxonomy or an analysis of species overlap between habitats and sites, while the second manuscript concentrates on providing species names and descriptions thus targeting taxonomic experts. To ensure that the information in both manuscripts is available at the same time, we recommend joint publication as preprints. It should be duly noted, however, that this requires the registration of all scientific names in Zoobank and the inclusion of a disclaimer according to section 8.2. of the ICZN code that the preprint is not issued for the purpose of zoological nomenclature and should be considered not published within the meaning of the code.

In this paper, we propose and test “dark taxonomy” by delimiting the species and describing the fauna of Mycetophilidae of Singapore based on an initial batch of 1,454 specimens. We then test the efficiency of the protocol for discovering the most abundant species by using a second batch of 1,493 specimens to measure the proportion of species already described based on the first batch. In addition, we illustrate the size of the taxonomic impediment for dark taxa in the Oriental region by demonstrating that revising the fauna of a small island increases the number of described species by >25%.

## Material and Methods

The project proceeded through five phases, but the first phase (collecting) generated not only samples for this study, but also for numerous others revising different dark taxa; i.e., many dark taxonomy studies start with phase 2.

### Phase 1: Collecting

Malaise traps were deployed for varying periods (2-6 months) between April 2012 and June 2019 at 107 sites in Singapore (Figs. 1A–B, see details in Amorim et al., 2023 and Yeo et al., 2021).The sites represented the following habitat types in Singapore: degraded urban secondary forest, old and maturing secondary forest, primary forest, coastal forest, swamp forest, freshwater swamp (lacking mature trees), and old-growth and replanted mangrove forests (Fig. 2A– E). The samples were collected weekly and preserved in 70% ethanol. Subsequently, they were sorted to order/family by parataxonomists and entomologists. Mycetophilids were present in most sites, though not particularly abundant in some habitats (e.g., mangroves).

**Figure 1.**
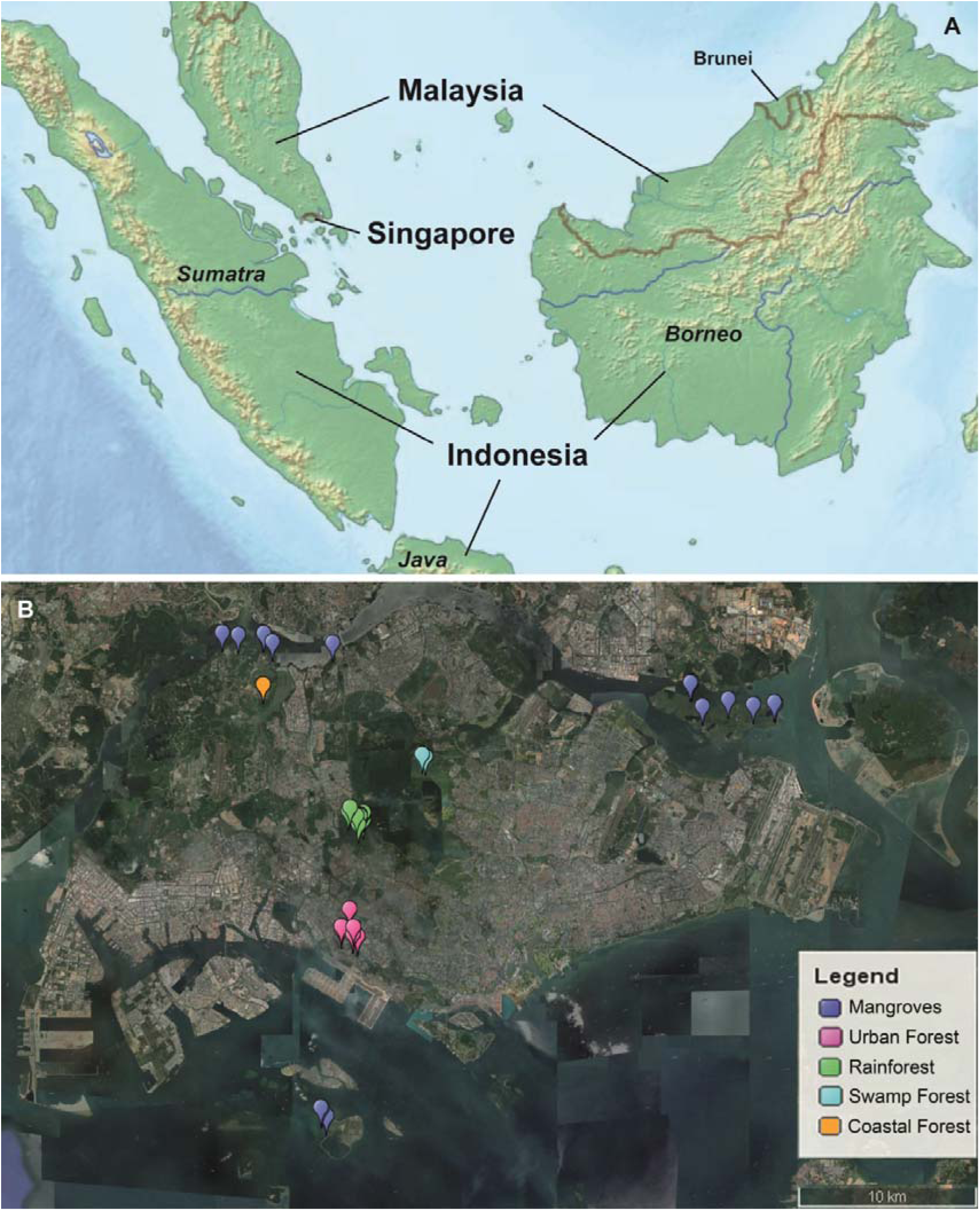
Singapore sites sampled with Malaise traps (modified from www.freeworldmaps.net). A. Southeast Asia, with relative position of Singapore. B. Collecting sites in different environments.

**Figure 2.**
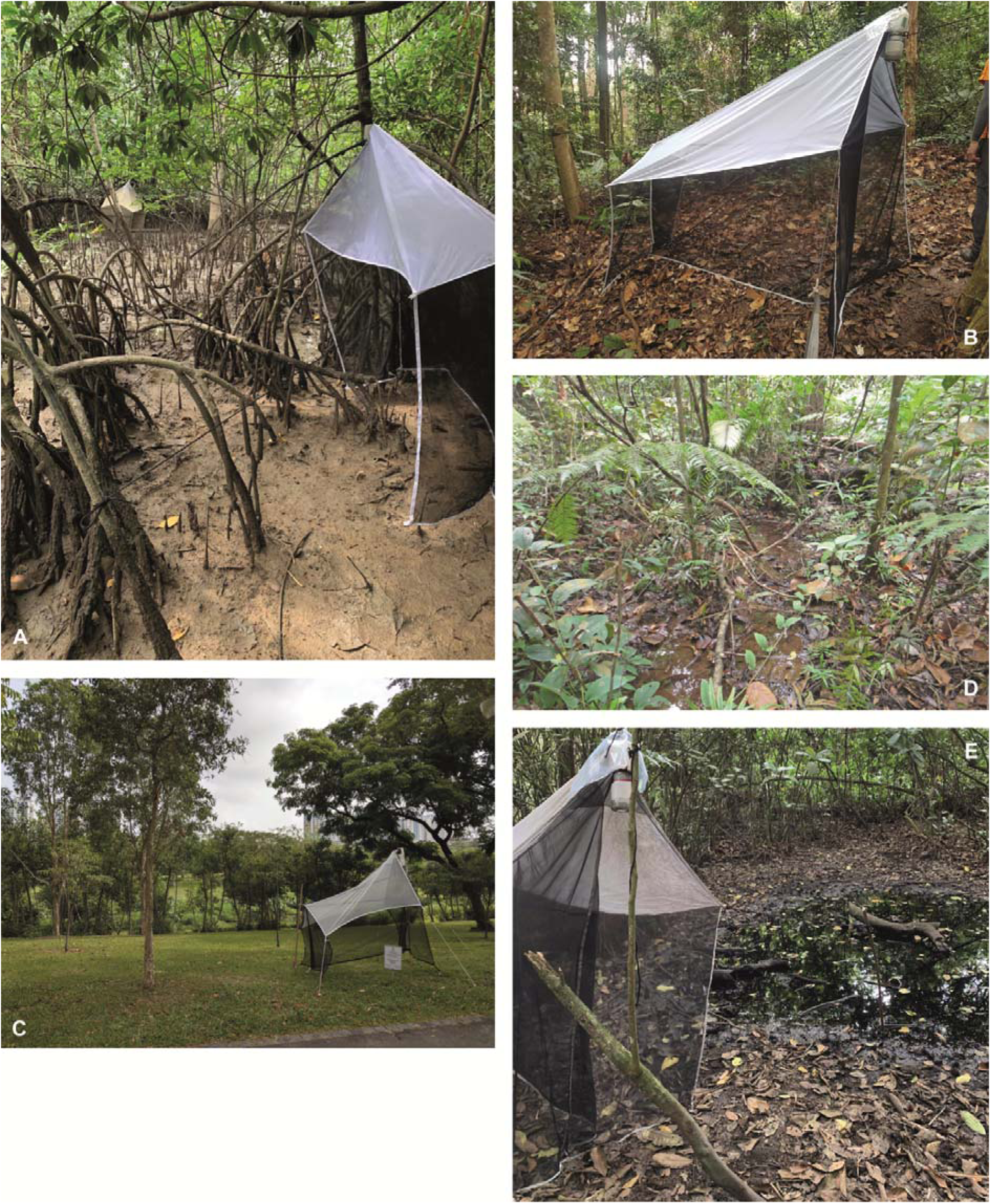
Environments sampled in Singapore. A: Mangrove, B: Rainforest, C: Urban forest, D: Swamp forest, E: Freshwater swamp.

### Phase 2: Estimating the number of MOTUs

We processed two batches of specimens. The first had 1,454 and the second, 1,493 specimens. Ideally, an assignment of the specimens to batches should have been done randomly, but the batches evolved according to the order in which the specimens were sequenced in the laboratory and the morphology of the sequenced specimens was studied. For the latter, there was a cut-off date because a taxonomic monograph had to be prepared (Amorim et al. 2023). We therefore later tested whether this haphazard versus random assignment to batches would have affected the overall results using randomized assignments (see below).

All specimens were sequenced for a 313-bp fragment of the *cytochrome oxidase I* gene (*COI*) since such minibarcodes amplify better than the full-length barcode (Srivathsan et al. 2021) and nevertheless yield similar species-level signal (Yeo et al. 2020). The minibarcode was amplified for each specimen using a protocol in Meier et al. (2016). In the early phases of the study, DirectPCR (Wong et al. 2014) was employed; i.e., 1 or 2 legs of each specimen were used as the DNA template for amplification. For specimens collected later, DNA extraction was performed by immersing the whole specimen in Lucigen’s QuickExtract solution or HotSHOT buffer (MonterolJPau et al. 2008). PCR was performed with the following primer pair mlCO1intF: 5’-GGWACWGGWTGAACWGTWTAYCCYCC-3’ (Leray et al. 2013) and jgHCO2198: 5’-TANACYTCNGGRTGNCCRAARAAYCA-3’ (Geller et al. 2013). These primers were labelled at the 5’ end with 9-bp tags that differed from each other by at least three base pairs. Every specimen in each sequencing library was assigned a unique combination of forward and reverse primer tags. This allows for reads to be assigned to their specimen of origin. A negative control was used for each 96-well PCR plate to detect contamination. Amplification success was estimated via gel electrophoresis of a subsample of eight wells from each plate.

Amplicons were pooled at equal volume within each plate. Equimolar pooling across plates was performed by approximating concentration based on the presence and intensity of the gel bands. The pooled samples were cleaned with Bioline SureClean Plus before being sent for library preparation at AITbiotech using TruSeq Nano DNA Library Preparation Kits (Illumina) or the Genome Institute of Singapore (GIS) using NEBNext DNA Library Preparation Kits (NEB). Paired-end sequencing was performed on Illumina Miseq (2x300-bp or 2x250-bp) or Hiseq 2500 platforms (2x250-bp). The sequences were obtained across multiple runs, which allowed for troubleshooting and re-sequencing PCR products which failed to yield a sufficiently high number of reads during the first sequencing attempt. Some of the specimens were also sequenced with MinION (Oxford Nanopore) using primers labelled with slightly longer tags (13-bp; see Srivathsan et al. 2019). The raw Illumina reads were processed with the bioinformatics and quality-control pipeline described in Meier et al. (2016). A BLAST search to NCBI GenBank’s nucleotide (nt) database was also conducted to identify contaminant sequences by parsing the BLAST output with *readsidentifier* (Srivathsan et al. 2015). Barcodes that matched non-target taxa at >97% identity were discarded.

Barcodes lacking stop codons were aligned with MAFFT v7 (Katoh and Standley 2013) using default parameters and analysed using three different species delimitation algorithms: objective clustering (Meier et al. 2006), Automatic Barcoding Gap Discovery (ABGD: Puillandre et al. 2012) and Poisson Tree Process (PTP: Zhang et al. 2013). Objective clustering was implemented via a python script that implements the objective clustering algorithm in Meier et al. (2006) and we obtained the results for four p-distance thresholds (2–5%) encompassing thresholds commonly used in the literature for species delimitation (Ratnasingham and Hebert 2013). ABGD was performed using uncorrected p-distances and the minimum slope parameter (-X) 0.1, with the default range of priors (P = 0.001 – 0.1). Lastly, a maximum likelihood phylogeny was generated from the barcode sequences in RAxML v.8 (Stamatakis 2014) with the GTRGAMMA model and with 20 independent tree searches. The barcode tree was then used for the PTP analysis using the implementation provided in mPTP (--ml --single) (Kapli et al. 2017; Zhang et al. 2013) algorithms. As described in Wang et al. (2019), the vials with the specimens were then physically sorted into bags that corresponded to 3% p-distance clusters. Specimens from these bags were used for morphological verification of species boundaries according to the rules of LIT.

### Phase 3: Integrative Taxonomy

We started the morphological study before LIT (Hartop et al., 2022) was published. We thus slide-mounted at least one male and one female (if available) for each 3% MOTU without picking specimens representing particular haplotypes. For slide-mounting, the specimens were cleared with KOH, dehydrated in ethanol, dissected to separate wings, abdomens with terminalia, and head/thorax. The body parts were mounted in Euparal (modified from Walker and Crosby 1988; Huber and Reis 2011) under three separate coverslips. Particularly, the specimens that underwent DNA extraction with Proteinase K overall worked well because the proteins were already digested, while the exoskeleton was well preserved for slide mounting (Santos et al. 2018). However, some soft specimens collapsed, affecting the quality of slide mounts.

This initial screen was then enhanced to also test the LIT rules outlined in Hartop et al. (2022). All non-singleton barcode clusters obtained at 3% were classified as stable or instable based on a stability index that quantifies whether cluster content changes when the clustering threshold is increased from 1% to 3%. The clusters where the maximum pairwise distance between any two specimens was >1.5% were also flagged as instable. All stable clusters were tested with morphology by studying a pair of specimens representing the most distant haplotypes, while for all instable clusters the morphology of one specimen representing the main haplotypes was also studied. For most genera, the diagnoses include male terminalia features, such that only clusters with males could provide sufficient information on conspecificity. For each cluster we then assessed whether (1) the examined specimens from the same MOTU belonged to the same morphospecies and (2) differed from the morphospecies in the “neighboring” clusters. If (1) was violated, the cluster was split to test whether morphospecies and DNA barcode information were congruent at a lower clustering threshold. If (2) was violated, it was tested whether fusing “neighboring” clusters restored congruence with morphospecies. Eventually, we accepted those grouping decisions based on DNA barcodes that were congruent with morphological information. No species delimitation was performed when morphospecies were incongruent with all grouping statements based on barcodes.

### Phase 4: Species identification and description

To decide whether a specimen belonged to a described species or one that had to be newly described, we extensively reviewed the literature on the Oriental fauna of Mycetophilidae, which is here also cited because it forms the foundation of all taxonomic work on the group (Meier 2017). For the Oriental species described by Edwards (1925, 1926, 1927, 1928, 1929, 1931, 1933, 1935, 1940), we studied and photographed primary types deposited at the Natural History Museum, London. For more recently described Oriental species of Mycetophilidae, we were able to resolve species identity based on published careful descriptions with good illustrations (Colless 1966; Zaitzev 1982; Sivec and Plassmann 1982; Wu and Yang 1986; Bechev 1995; Søli 1996, 2002; Kallweit 1998; Matile 1999; Xu and Wu 2002; Wu et al. 2003; Papp 2004; Hippa et al. 2005; Hippa 2006, 2007, 2008, 2009, 2011; Ševčík 2001; Ševčík and Hippa 2010; Hippa and Ševčík 2010, 2013; Ševčík et al. 2011, 2012; Borkent and Wheeler 2012; Ševčík & Kjærandsen 2012; Kurina and Hippa 2015; Hippa and Saigusa 2016; Hippa and Kurina 2018; Magnussen et al. 2019; Fitzgerald 2017; Kaspřák et al. 2017; Kjærandsen et al. 2023). However, some Oriental species, especially those from Sri Lanka, could not be resolved because the descriptions were insufficiently detailed, and the types were lost or unavailable for study. We here assumed that the species from Singapore are allospecific based on geography, which is consistent with high endemism for Mycetophilidae species in South America (Amorim and Santos 2018), although our revision (Amorim et al. 2023) found a species whose distribution ranges from Japan through Singapore and Sumatra to Thailand. Future study will have to reveal if any of the Sri Lankan species are conspecific with the material collected in Singapore.

### Phase 5: Completeness of the faunal assessment

The barcode data for the second batch of specimens (N=1,493) was used to test how complete the revision based on the first batch of specimens was (N=1,454). For this purpose, the genetic data were analyzed as described earlier for the first batch of sequences. Subsequently, we assessed how complete the initial revision was in the light of the species diversity represented in the second set of specimens. We also determined the number of new haplotypes, clusters, and how many specimens from batch 2 belonged to species described based on batch 1 or new species. We then compared these observed values to those obtained by randomized assignment of specimens to batch 1 and batch 2, assuming the same sample sizes (batch 1, N=1,454 and batch 2, N=1,493). Randomization was carried out using a custom script (https://github.com/asrivathsan/darktaxa). Lastly, we estimated how many specimens would require morphological study according to LIT to validate MOTUs for all 2,947 specimens in both batches.

## Results

### Phase 1: Collecting

The material for this study was obtained during 3,526 Malaise trapping weeks distributed across five different habitats in Singapore: mangroves (74 sites, 2,162 trap weeks), tropical forest (nine sites, 567 weeks), urban forests (15 sites, 280 weeks), swamp forest (four sites, 262 weeks), coastal forest (10 sites, 156 weeks), and freshwater swamp (seven sites, 99 weeks). A total of 69 traps collected 3,030 mycetophilid specimens that were successfully sequenced. Note that therefore the overall number of mycetophilid specimens collected was somewhat lower than expected for such a large number of samples. This is likely due to fact that almost two thirds of the samples (61%) were obtained from mangroves that are comparatively species-poor for Mycetophilidae.

### Phase 2: Molecular processing

The 1,454 COI barcodes from the first batch clustered into 115–128 MOTUs via objective clustering at 2–5% uncorrected p-distances (Table 1). With ABGD, the number of MOTUs ranged from 115–128. PTP clustered the barcodes into 127 MOTUs. The 1,493 COI barcodes from the second batch yielded 95 MOTUs from 5% p-distance clustering.

**Table 1.** Congruence and conflict between molecular and morphological data for batch 1 using three different algorithms for MOTU estimation (PTP, Objective Clustering, ABGD). For each algorithm and parameter, the table indicates the number of morphospecies that were congruent with the respective MOTUs, the number of morphospecies that were split into more than one species, and the number of morphospecies that were lumped. Note that there were 117 morphospecies with molecular data that could be used to determine the match ratios.

| Algorithm | PTP | Objective Clustering |  |  |  | ABGD |  |  |  |  |
| --- | --- | --- | --- | --- | --- | --- | --- | --- | --- | --- |
|  |  | 2% | 3% | 4% | 5% | P=0.001<br>-0.005 | P=0.008 | P=0.013<br>-0.022 | P=0.036 | P=0.06<br>0 |
| No. MOTUs | 127 | 128 | 124 | 121 | 115 | 126 | 125 | 124 | 122 | 115 |
| No. congruent morphospecies | 108 | 107 | 110 | 113 | 113 | 109 | 110 | 110 | 112 | 113 |
| Split morphospecies | 9 | 10 | 7 | 4 | 0 | 8 | 7 | 7 | 5 | 0 |
| Fused morphospecies | 0 | 0 | 0 | 0 | 2 | 0 | 0 | 0 | 0 | 2 |
| Match ratio | 89% | 87% | 91% | 95% | 97% | 90% | 91% | 91% | 94% | 97% |

### Phase 3: Integrative Taxonomy

Congruence between morphospecies and MOTUs was quantified using the match ratios (match ratio = 2∗N_congruent_/(N_MOTU_+N_morph_); Ahrens et al., 2016). It was overall high (0.87-0.97), with the highest match ratio observed for 5% clusters and MOTUs obtained with ABGD at P=0.06 (0.97, 113 morphospecies congruent).

### Phase 4: Species identification, description and naming

The application of LIT to the 124 MOTUs obtained with a 3% clustering threshold required morphological study of 224 specimens. Of these 124 MOTUs, 27 were singletons, 73 MOTUs were stable (937 specimens), and 24 were instable. One challenge with carrying out the LIT analysis was that for approx. 40% of haplotypes, only female specimens were available. This weakens the congruence test between morphology and molecular data, because females have fewer informative morphological characters. Overall, we find that obtaining MOTUs congruent with morphospecies required fusing 3% MOTUs in seven cases (Figure 3); i.e., we were able to delimit 117 species with morphology and molecular data. They belonged to 21 known and one new genus (note that the monograph also describes three additional species based on morphology only: see details in Amorim et al. 2023). Only five species were already described prior to this dark taxonomy study: *Chalastonepsia hokkaidensis* Kallweit, *Eumanota racola* Søli, *Metanepsia malaysiana* Kallweit, *Parempheriella defectiva* Edwards, *Neoempheria dizonalis* Edwards. The remaining are described and illustrated in the companion taxonomic monograph (Amorim et al. 2023). Images are also available from digital reference collection “Biodiversity of Singapore” (https://singapore.biodiversity.online/).

**Figure 3.**
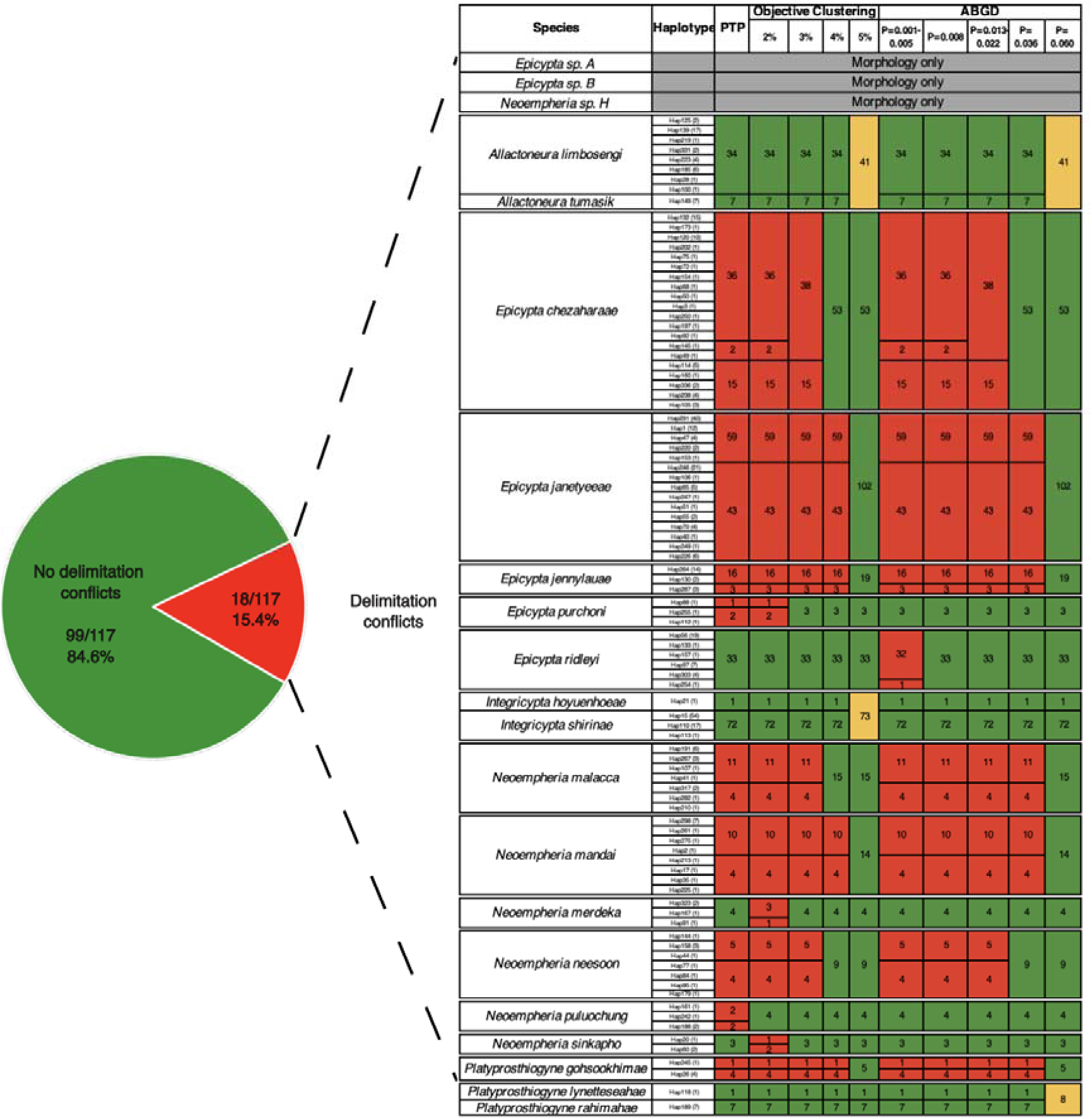
Most MOTUs are stable and congruent with morphospecies (green; N=99). Remaining species have splitting (red) and lumping (yellow) conflict with molecular data depending on species delimitation algorithm.

### Phase 5: Completeness of the faunal assessment

The second batch of specimens yielded 94 MOTUs at 5% in a combined analysis of all the 2,947 barcodes from both batches. 5% MOTUs were here used for clustering because it maximized congruence between morphology and molecular data for batch 1. Of these, 76 belonged to MOTUs already examined for species descriptions based on the specimens in the first batch (76/94 or 80% of species in batch 2). All 18 newly discovered MOTUs were stable (no compositional changes with all delimitation algorithms and settings used here) and were so rare that they only represented 31 of the 1,493 specimens in the second batch; i.e., 97.9% of all specimens in batch 2 belonged to MOTUs that were also found in batch 1 (Figure 4). To assess if the coverage of species and specimens for batch 2 was influenced by sampling, we compared the observed values in batch 1 and batch 2 to those obtained through randomized assignments of specimens to either batch (100 randomizations). We find that randomized assignments yield similar values as the observed ones (observed MOTUs: 94, randomized MOTUs: 82 ± 2.7%; observed coverage: 97.9%, randomized coverage: 98.1 ± 0.4% (Figure 4).

**Figure 4.**
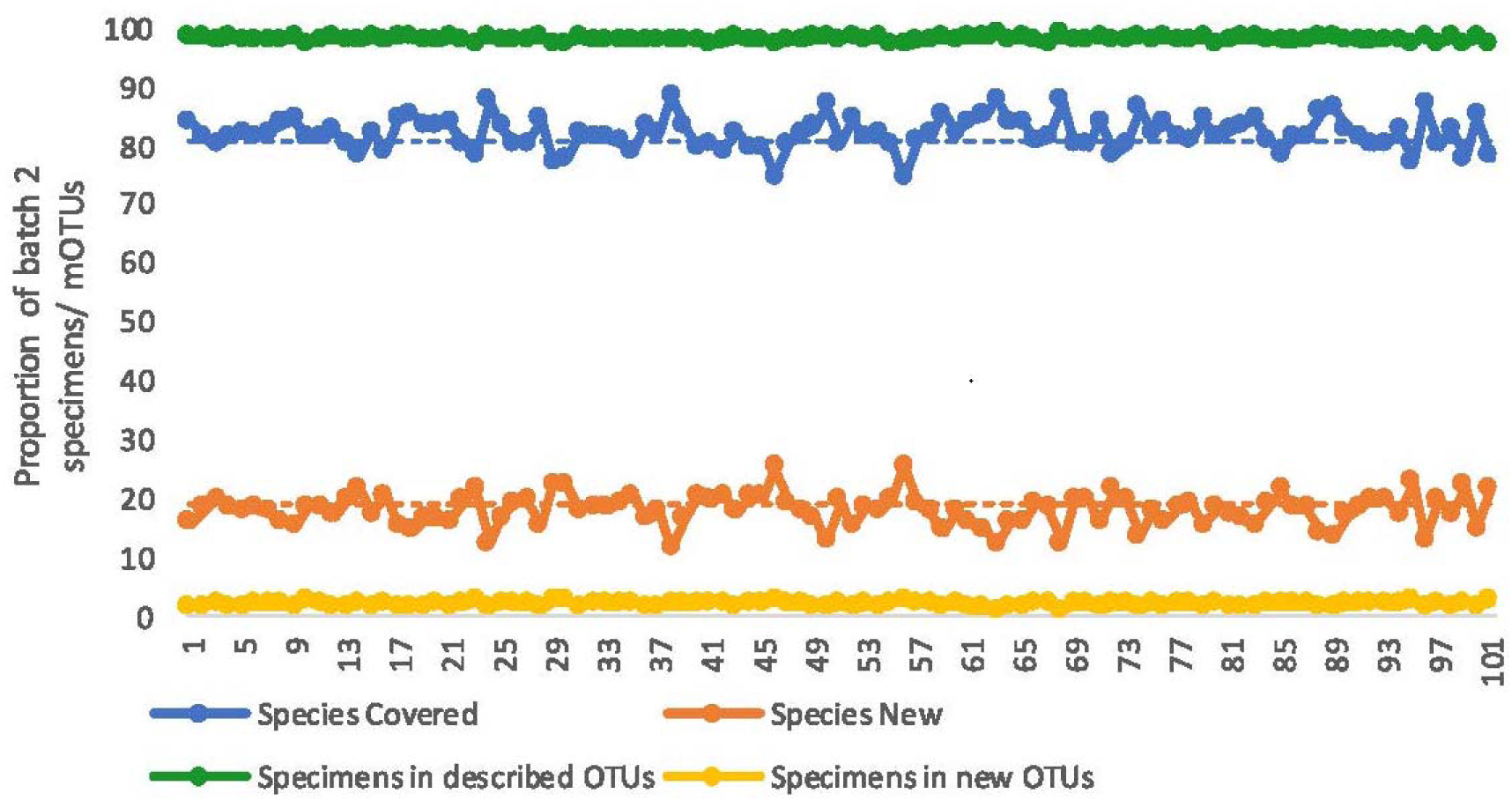
An overwhelming number of specimens and most species in specimen batch 2 belong to species described based on specimens in batch 1 (∼80% of species and >97% of specimens). Dashed lines represent the observed values while the solid lines represent the results of 100 randomizations (x-axis).

At 5% objective clustering, the proportion of singletons was 22% (N=29) for all specimens, 17% (N=25) for dataset 1, and 26% (N=24) for dataset 2. The corresponding numbers for doubletons were 11% (N=14), 9% (N=11), and 16% (N=15). Carrying out a LIT analysis for all 2,947 specimens would have required checking 143 (3%) MOTUs of which 30 were singletons, 79 stable, and 34 were instable (1,741 specimens). Despite the large number of specimens in the instable clusters, only 272 specimens of the combined dataset would have to be checked to sample haplotypes according to LIT.

## Discussion

Biodiversity loss is now considered one of the top five threats to planetary health (World Economic Forum 2023). Yet, most animal species are undescribed, unidentifiable, and lack baseline data for biomonitoring. Much of this diversity apparently belongs to dark taxa such as the top 20 species-rich families of flying insects that were identified in Srivathsan et al. (2023) based on samples from five biogeographical regions. For many of these taxa, we know how to obtain standardized samples (e.g., Malaise, pitfall, flight intercept traps), but most samples are currently only evaluated with metabarcoding (Yu et al., 2012). However, bulk sequencing is unable to overcome taxonomic impediments because it disrupts the association between specimen and genetic data. In addition, metabarcoding only provides approximate abundance/biomass data. However, even those specimens that are subjected to high-throughput barcoding (“megabarcoding: Chua et al., 2023) tend to remain in the taxonomic desert because they are not morphologically identified beyond family level. For example, in 2021 95% of all Sciaroidea and 60% of all Mycetophilidae specimens were found to lack species-level identifications (Kjærandsen 2022). To overcome such problems, we propose that the principles of “dark taxonomy” should be used to delimit species, obtain species-specific barcode libraries, and facilitate taxonomic documentation through description and naming for the species in at least some samples at each biomonitoring site. This will also generate biodiversity baselines that include abundance information.

Our study illustrates how this is feasible even for dark taxa. We show that dark taxonomy can fairly quickly yield a good estimate of species diversity and provide species names and barcodes for most common species belonging to a dark taxon. Our initial set of specimens consisted only of 1,454 specimens. They are shown in the companion monograph (Amorim et al., 2023) to belong to 117 species that could be delimited with morphological and molecular data. We then tested the completeness of the revision by using the data for an additional 1,493 specimens. Fortunately, the observed number of MOTUs increased by only 16% (N=18); i.e., the richness estimate based on batch 1 was more than 80% complete. Furthermore, 76 of the 117 species in the first batch were also represented in the second batch. Not surprisingly, they were mostly common species, so that the proportion of specimens from the second batch that belong to species described in the first batch is very high (>97%). This means that one dark taxonomy study was here sufficient for readying the fauna of one dark taxon for a small country like Singapore-. Most mycetophilid species will now be identifiable based on molecular or morphological data thus facilitating the inclusion of this taxon in biomonitoring. This is impressive given that the number of known and described species of Mycetophilidae alone in Singapore now exceeds the number of species in major vertebrate groups (mammals: 83 spp.; amphibians: 26 spp.; “reptiles”: 109 spp; source: Singapore National Parks Board [Red Data Book] Species List, 2023). Moreover, all the new species can be sampled with Malaise traps, which have minimal environmental impact.

Currently, very little is known about the natural history of tropical fungus gnats so it is difficult to assess whether they could be used as indicator taxa for habitat quality or fungus richness. However, this seems likely, given studies such as the ones by Økland (1994, 1996) demonstrating that mycetophilids are good indicators of temperate forest quality. However, more environmental correlates need to be collected for the tropics and it will be important to associate larvae and adults (see Yeo et al. 2018) to understand to what extent mycetophilid species are specializing on specific species of fungi and how mycetophilid species-richness and abundance could be used as indirect evidence of tropical forest health.

To fully appreciate the potential of dark taxonomy, it is important to consider that Srivathsan et al. (2023) showed that more than half of the specimens and species in Malaise trap samples belong to approximately 20 family-level dark taxa (see also Brown 2005; Karlsson et al. 2020). This means that applying a technique like “dark taxonomy” to a few complete Malaise trap samples could simultaneously yield biodiversity baseline data for many numerically dominant families and hundreds of species. This is realistic, because barcoding is now so easy and cheap (Meier, et al., 2024; Srivathsan et al. 2021; Srivathsan et al., 2024) that dark taxonomy is a frugal techniques and thus suitable for biodiverse countries with limited science funding (Brydegaard et al. 2024). PCR costs can be as low as 0.05 USD per specimen, and sequencing with Illumina’s NovaSeq costs only 0.01-0.02 USD /barcode (Srivathsan et al. 2021). A self-contained barcoding lab covering all steps from specimens to barcodes can be installed for <USD 10,000 if it utilizes Oxford Nanopore Technologies (ONT) sequencers (e.g., MinION; Srivathsan et al. 2024). The use of 3^rd^ generation sequencing technologies raises the sequencing cost for DNA barcodes, but it still means that the consumable cost for barcoding the ∼3,000 specimens in our study costs USD 150 (Illumina) to USD 375 (ONT). Specimen handling and imaging costs are also dropping with the use of DIY microscopes (Wührl et al. 2024) and robots for sample sorting (see Wührl et al. 2022). Carrying out taxonomic revisions for dark taxa will inevitably remain challenging but “dark taxonomy” can help with simultaneously addressing taxonomic impediments and the need for biodiversity baseline data.

### Objections to dark taxonomy

The proposal of “dark taxonomy” is likely to be controversial for a variety of reasons. One is that specimens/types from dry collections will initially only be used if they are important for resolving species names. This means that many rare and all extinct species will not be covered by dark taxonomy. However, we would argue that the advantages of dark taxonomy outweigh these disadvantages. Concentrating on fresh material from standardized traps means that specimens are suitable for barcoding. Secondly, standardized samples come with complete metadata and there is no need to digitize thousands of specimen labels individually. None of this can be said for taxonomic revisions that deal with a mixture of wet and dry specimens. This does not mean that dry material will become irrelevant. Dark taxonomy is designed to rapidly improve taxonomic knowledge for a poorly known taxon to reach the level where conventional taxonomic methods can be applied. At this stage, pinned museum specimens will become critical for testing uncertain species boundaries and estimating the full species-level diversity of a clade.

We also suspect that some systematists may be concerned that dark taxonomy proposes to work at small geographic scales. However, there are two reasons why we are less worried. Firstly, we would like to point to the success of those turbo-taxonomic studies that seem to have led to few cases, if any, where species boundaries had to be revisited later when the fauna of neighboring areas was covered. The best example is the weevil genus *Trigonopterus*, where several hundred species have been described in regional treatments (Narakusumo et al. 2019, 2020; Riedel 2022; Riedel and Narakusumo 2019; Riedel and Tänzler 2016; Riedel et al. 2013b, 2014; Van Dam et al. 2016). Of course, only time will tell as to which proportion of species proposed based on poor geographic sampling will have to be revised as geographic sampling increases, but it appears unlikely that the proportion will be high as long as delimitation is based on molecular and morphological data. Our prediction is based on the current observation that barcodes alone tend to successfully delimit the majority of species. Note, however, that the precise proportion remains unknown given that most species still lack barcodes and comparatively few species have been sampled across their full geographic range. Denser sampling tends to obscure the signal in barcode data as can also be seen in our study given that the proportion of specimens in instable MOTUs increased from 530 in batch 1 (36%) to 1,741 in the combined batches (59%). However, overall, we remain optimistic that the vast majority of species delimited with dark taxonomy will be stable, because the species boundaries were determined by morphological and molecular data. So, both data sources would have to be misleading to generate incorrect species limits.

Those systematists who are still reluctant to accept dark taxonomy may want to consider the alternative. By describing more than 100 species based on 1,454 specimens collected in a small country such as Singapore (730 km^²^), we are increasing the number of known Oriental species by >25%. This indicates an alarming neglect and taxonomic impediment given that Singapore is expected to have a comparatively low species diversity because it has lost most of its natural habitats, and many species belonging to charismatic taxa are known to have gone extinct (Chisholm et al. 2023; Theng et al. 2020). Expanding the geographic scale of a mycetophilid revision to the Malay Peninsula would mean revising the fauna of an area that is 300 times larger (242,363 km^2^), with more pristine habitats and elevational gradients that are likely to increase the species diversity. It would surely be impossible to handle the fauna at this scale using conventional means. One may propose to tackle the fauna one genus at a time, but this would still not be feasible for three species-rich genera that contribute over 60% of the new species described in the companion monograph (*Neoempheria* Osten-Sacken: N=31; *Epicypta* Winnertz: N=29; *Manota* Williston: N=14). Overall, we would thus argue that the alternative to dark taxonomy is dangerous neglect given that dark taxa contain so many species.

Of course, working at small geographic scales will require regular combining of information from multiple faunistic treatments covering neighboring areas. For example, our dark taxonomy study of Singapore’s Mycetophilidae will become essential background information for a taxonomic revision of the same group in additional studies on the fauna of the Malay Peninsula. Fortunately, combining and analyzing barcode data from different studies is straightforward (Meier et al. 2021; Vences et al. 2021) as long as the same barcoding region is sequenced. Such analysis then yields information on the stability and intraspecific variability of each MOTU. This information furthermore guides an expanded LIT analysis involving morphological and/or nuclear data. It is important that this follow-up work should not require the loan of type material to avoid delays and the cost and risks of specimen shipping. This is one major reason why the morphology of new species should be well documented (Ang et al. 2013). But such documentation is also important for other reasons. It is the only way to provide comparison data for species that were described based on morphology over the last 250 years. This is almost all, given that even now only 10% of all new species descriptions in entomology contain molecular data (Miralles et al. 2021). Integrative species descriptions are also critical for the analysis of specimens not suitable for sequencing and for allowing researchers without access to molecular labs to participate in taxonomy, which constitutes a large number of biologists all over the world (Zamani et al. 2022a,b). Neglect of morphology also seems very shortsighted, given that it appears likely that specimen identification with molecular tools will eventually be complemented with identification based on image-based AI tools (Høye et al. 2021, Wührl et al. 2022, 2024). These tools need to be trained with images and arguably the best images come from taxonomic revisions because the specimens have been competently identified (Meier and Dikow 2004).

### Establishing integrative species limits with LIT

An important aspect of our study was testing the performance of Hartop et al.’s (2022) Large-Scale-Integrative Taxonomy (LIT). We find that it performs well for Mycetophilidae. Although the initial clustering of barcodes/specimens was at a threshold of 3% that was later found to be too low, the number of specimens that had to be studied to identify MOTUs that were congruent with morphospecies was moderate (N=224). The proportion of specimens was higher than in the previous application of LIT (Mycetophilidae: 15.4%; Phoridae: 5.1% in Hartop et al. 2022), but this was largely due to the high species/specimen ratio of 8% in Mycetophilidae. Carrying out a LIT analysis for all data (N=2,947) would reduce the proportion of specimens to be studied to 9% (Table 2). In the case of Singaporés Mycetophilidae, the level of congruence between MOTUs used for specimen sorting (3% clusters) and morphological data was very high (110 congruent MOTUs). This means that only seven morphospecies were in conflict. Note also that the term “conflict” is hardly appropriate from an evolutionary point of view. Measurements such as the match ratio only quantify which proportion of morphospecies are in perfect agreement (i.e., form identical sets) with MOTUs obtained using one particular clustering threshold or algorithm. Yet, the use of a single threshold for clustering barcodes has long been criticized as inappropriate on theoretical grounds (Will and Rubinoff, 2004). Indeed, we would expect that closely related species should form several proper subsets within a single MOTU if the latter was delimited using a threshold appropriate for “average species”. Conversely, we would expect old species to form union sets composed of several MOTUs. These expectations are in line with the results of a recent empirical study on optimal sequence similarity thresholds (e.g., Bonin et al. 2023). LIT therefore not only establishes whether morphospecies are “in conflict” with MOTUs, but also tests whether there are clustering thresholds that yield congruent MOTUs. Only if this is not the case, LIT would consider barcode and morphological data to be in genuine conflict and the species would remain undescribed until additional data become available (e.g., nuclear markers, ecology). However, it is comforting that we did not find such a case in our current study or our recent study on Swedish phorids (Hartop et al. 2022).

**Table 2.** Comparison of the number of MOTU (3%) and specimens in the first batch and all data.

| Dataset | Batch 1 | All data<br>(batch 1 + 2) |
| --- | --- | --- |
| 3% MOTUs (No. of specimens) | 124 (1454) | 143 (2947) |
| Singleton MOTUs | 27 | 30 |
| Stable MOTUs (No. of specimens) | 73 (897) | 79 (1176) |
| Unstable MOTUs (No. of specimens) | 24 (530) | 34 (1741) |
| LIT: No. of specimens to be examined<br>(singleton/stable/unstable) | 224 (27/130/67) | 272 (30/136/106) |

### Publishing dark taxonomy revisions

Many taxonomic monographs consist of two main sections. The introduction and discussion sections contain information that is of interest to a broad readership (e.g., information on biology, species diversity, taxonomic history of a taxon) while the taxonomic section consists of species descriptions that are mostly of interest to taxonomic specialists. Unfortunately, the information that is of general interest is often overlooked by biologists and thus also not cited. We here suggest that a dark taxonomy revision should be published in two separate publications. The taxonomic monograph could only consist of an introduction/method section that presents the background information needed to understand how the species hypotheses were derived, as well as the taxonomic descriptions. Everything else could be in a separate companion paper intended for a broader audience. Such a companion manuscript can, for example, present a rigorous quantitative analysis of the habitat preferences and phenology of species if the revision was based on fresh samples obtained from standardized samples. We believe that it is desirable to bring both manuscripts together in the form of simultaneous releases as preprints and this was the approach pursued here. In our general paper, we introduce the concept of dark taxonomy, demonstrate that a single, moderately sized revision can increase the number of described species in a dark taxon by >25% for the Oriental region, and present evidence that the species discovered in a first revision cover a very large proportion of specimens collected at different times. In the companion monograph, we describe and illustrate the species and discuss the taxonomic affinities of the newly described species (Amorim et al. 2023).

## Conclusions

Dark taxa are everywhere. They are abundant, species-rich, and a major obstacle to holistic biomonitoring. What is needed is a robust “dark taxonomy” protocol designed to tackle the taxonomic impediments for this dark diversity. We would argue that whatever protocol may eventually be used should satisfy two criteria: it should be efficient enough to deal with thousands of specimens and species and it should also yield biodiversity baseline data, so that changes in abundance and richness of dark taxa can be monitored in a not-too-distant future and utilized for ecosystem management or rehabilitation purposes. We believe that our proposal of “dark taxonomy” can be a major step in this direction. Robots and cost-effective sequencing enable presorting of species to putative species. LIT reduces the amount of morphological work needed to obtain integrative species boundaries. Lastly, barcoding of type specimens will accelerate the process of distinguishing between species that need a scientific name and those that are already described.

## Data availability statement

Data files and/or online-only appendices can be found in the Dryad data repository: https://datadryad.org/stash/share/xFTVi0kZueBDEU4AXU-xaNeNa9o-y_jYvSRFlQdjt7s. The custom script used for randomization of the data set partitions is available from https://github.com/asrivathsan/darktaxa.

## Acknowledgements

All work described here was carried out as part of a comprehensive insect survey of Singapore which was carried out in collaboration and with support from the National Parks Board of Singapore (NParks). Special thanks go to Dr. Patrick Grootaert and the team from the National Biodiversity Centre of NParks for their assistance in fieldwork (Permits: NP/RP12-022-4, NP/RP12-022-5, NP/RP12-022-6). We would also like to thank the research staff, lab technicians, undergraduate students, and interns from the National University of Singapore’s Evolutionary Biology Laboratory and Lee Kong Chian Natural History Museum for their help and assistance. This project would have been impossible without their hard work. Financial support was provided by Tier 2 funding from the Ministry of Education (Singapore) and the LKCNHM Research Visitor Fund.

